# Transcriptomic profiling of gill biopsies to define predictive markers for seawater survival in farmed Atlantic salmon

**DOI:** 10.1101/2024.08.20.608748

**Authors:** Lars Grønvold, Mattis J. van Dalum, Anja Striberny, Domniki Manousi, Trine Ytrestøyl, Turid Mørkøre, Solomon Boison, Bjarne Gjerde, Even Jørgensen, Simen R. Sandve, David G. Hazlerigg

## Abstract

Wild Atlantic salmon migrate to sea following completion of a developmental process known as parr - smolt transformation (PST), which establishes a seawater (SW) tolerant phenotype. Effective imitation of this aspect of anadromous life-history is a crucial aspect of commercial salmon production, with current industry practice being marred by significant losses during transition from the freshwater (FW) to SW phase of production. The natural photoperiodic control of PST can be mimicked by exposing farmed juvenile fish to a reduced duration photoperiod for at least 6 weeks before increasing the photoperiod in the last 1 - 2 months before SW transfer. While it is known that variations in this general protocol affect subsequent SW performance, there is no uniformly accepted industry standard; moreover, reliable prediction of SW performance from fish attributes in the FW phase remains a major challenge. Here we describe an experiment in which we took gill biopsies 1 week prior to SW transfer from 3000 individually tagged fish raised on 3 different photoperiod regimes during the FW phase. Biopsies were subjected to RNA profiling by Illumina sequencing, while individual fish growth and survival was monitored over 300 days in a SW cage environment, run as a common garden experiment. Using a random forest machine learning algorithm, we developed gene expression-based predictive models for initial survival and stunted growth in SW. Stunted growth phenotypes could not be predicted based on gill transcriptomes, but survival the first 40 days in SW could be predicted with moderate accuracy. While several previously identified marker genes contribute to this model, a surprisingly low weighting is ascribed to sodium potassium ATPase subunit genes, contradicting advocacy for their use as SW readiness markers. However, genes with photoperiod-history sensitive regulation were highly enriched among the genes with highest importance in the prediction model. This work opens new avenues for understanding and exploiting developmental changes in gill physiology during smolt development.

## Introduction

In anadromous salmonids, the transformation of freshwater resident juvenile fish (parr) into a migratory form (smolt) which goes to sea is known as smoltification, smolting or parr-smolt transformation (hereafter, PST). PST entails a complex combination of physiological and behavioural changes, amongst which the acquisition of the ability to efficiently maintain water and ionic balance is critical (McCormick, 2013).

PST typically occurs after several years of slow growth in freshwater (FW) streams where spawning took place and requires parr to have exceeded a certain minimum threshold size at the end of the growing season preceding PST (Handeland *et al*., 2013; Sigholt *et al*., 1995; Strand *et al*., 2018; Thorpe, 1977). Exposure to a declining autumn photoperiod followed by increasing photoperiod the following spring triggers the process of PST, apparently through a temporally coordinated sequence of endocrine change (McCormick, 2013). The photoperiod-dependent seasonal gating of PST ensures a synchronous springtime migration to sea, which is thought to reduce predation losses(Furey *et al*., 2016; Handeland *et al*., 1996).

Aquacultural production of Atlantic salmon (*Salmo salar*) depends on the ability to mimic natural PST to produce large numbers of seawater (SW) tolerant juvenile fish for transfer to sea cages where they can grow rapidly. There is no single universally accepted protocol for this commercial process. A widely used strategy for artificial smolt production is to rear hatchlings on a fast growth regime (i.e. continuous light (LL) and a fast growth diet for 6 - 12 months) to rapidly achieve threshold size, and then to expose them to short photoperiod (SP) for several weeks and before finally returning them to LL for the last 1 - 2 months of the FW phase. Based on observations of SW performance, it has been shown that the duration of exposure to SP should be at least six weeks long for LL to induce smolting (Duncan & Bromage, 1998). The mechanisms through which PST runs as a photoperiodic history-dependent process remain unknown, and untangling the role of SP exposure in smolt development is of considerable practical interest, as SP exposure reduces growth rates and slows aquaculture production (Imsland *et al*., 2014; Sigholt *et al*., 1998).

To overcome this delaying effect of SP exposure on production schedules, alternative strategies have emerged in which fish are maintained on constant light throughout the FW phase, and SW tolerance is achieved through dietary manipulation with salt / tryptophan enriched diets, water salinity manipulation, or by increasing size of fish (Striberny *et al*., 2021) in the run up to SW transfer. At present there is no unifying consensus on the relative merits of these different approaches, and difficulties in determining SW-readiness under different protocols probably contribute to industry losses during the SW-transfer phase of salmon production.

Because of its pivotal role in the regulation of water and ionic fluxes, undergoing a dramatic switch from a salt conserving (FW) to salt excreting (SW) function, the gill has become the focus of efforts to optimise commercial smolt production (Takvam *et al*., 2024). Within the gill, mitochondria-rich cells (MRCs) are considered the primary drivers of ionic regulation (Evans, 2008), and PST includes a pronounced shift in the location and phenotypic attributes of MRCs in the gill (McCormick, 2013; West *et al*., 2021). During PST, the gill complement of MRCs shifts from an ion-absorbing FW type to an ion-secreting SW type, and the distribution of MRCs shifts from the lamellae to the gill filament itself (Madsen *et al*., 2009; Pisam *et al*., 1988). One of the most studied molecular changes in a the gill MRC during PST is the shift in expression of the genes encoding the ion regulatory Na+, K+-ATPase (NKA) pump and concurrent increase in NKA protein activity (Reviewed in Takvam *et al*., 2024). Associated with this, relative levels of expression of two isoforms of the NKA α-subunit shifts during PST, with α1a showing higher expression in parr and α1b showing higher expression in smolts (McCormick *et al*., 2009).

From an applied perspective, characterisation of gill transcriptomic changes during PST provides a potential route to designing marker-based predictive strategies for optimising smolt production. Current approaches focus on relative expression of NKA alpha subunit isoforms (Takvam *et al*., 2024), but there is clearly potential to exploit other aspects of transcriptomic change to optimise predictive power. We have previously hypothesised that genes with photoperiod-history dependent regulation in the developing smolt gill would be interesting candidates for novel smolt status markers (Iversen *et al*., 2020). In this study we aim to test this, using a large-scale experiment in which gene expression prior to entering SW is compared with individual fish performance in the SW phase, including survival after SW transfer and stunted growth (i.e. loser fish (Noble *et al*., 2018)). We raised 3000 tagged fish on 3 different light regimes, produced RNAseq from gill biopsies one week prior to SW transfer, and subsequently recorded individual survival and growth in SW. We then applied a machine learning approach to this unique dataset, attempting to link SW performance to FW gill gene expression profiles. Our results demonstrate the potential for the development of novel and improved markers for smolt status in salmon aquaculture.

## Materials and Methods

### Fish rearing experiment

For detailed description of the fish rearing see Gjerde et al. (2024). In brief, three groups of about 1000 juvenile Atlantic salmon (*Salmo salar*) were exposed to three different photoperiods; LL (24 hours light), 12:12 (12 hours light / 24 hours) and 8:16 (8 hours light / 24 hours). The LL group was kept from hatching until sea water (SW) transfer on continuous light. The 8:16 group was reared on LL from hatching until the fish reached a body mass of ∼50g after which they were exposed to a 6-week period of short photoperiod exposure (8L:16D - 8 hours day light and 16 hours darkness). After these 6 weeks followed another 8 weeks on LL before sea water transfer when the fish reached a body mass of ∼100g. The 12:12 group was, exposed to 12h day light and 12h darkness for 6 weeks, following 8 weeks in LL before SW transfer. One week prior to sea water transfer, gill biopsies were sampled and smolt index (i.e. skin colouration), body weight and length were measured. The smolt index score is a categorical scoring of skin colouration; parr marks (score = 1), a mix of silvery skin with residual weak parr marks (score = 2), and completely silvery skin (score = 3).

Approximately one week after gill biopsies were taken, the fish were moved to one common seawater cage where they were kept for 10 months and daily recording of mortality were carried out.

### Ethical statement

The experiment was performed according to EU regulations concerning the protection of experimental animals (Directive 2010/63/EU). Appropriate measures were taken to minimize pain and discomfort. The experiment was approved by the Norwegian Food and Safety Authority (FOTS id. number 25658).

### RNAseq data generation

Gill biopsies were flash frozen on dry ice and sent to QIAGEN (Germany) for RNA-isolation using the RNeasy Fibrous Tissue Mini kit. Isolated RNA samples were then shipped to Novogene (UK) for RNA-sequencing library construction which were sequenced using 2*150 bp pair end illumina sequencing.

### RNAseq read mapping

Prior to transcript quantification, adaptor sequences were trimmed off using fastp version 0.23.2 (Chen *et al*., 2018). Read mapping and transcript quantification was done using Salmon version 1.1.0 (Patro *et al*., 2017) with the Atlantic salmon transcriptome annotation from ENSEMBL from the Ssal_v3.1 genome assembly (Ssal_v3.1, GCA_905237065.2) in the ‘selective mapping-mode’. A transcriptome index file was used as decoy sequence to avoid the incorrect mapping of reads between annotated and unannotated yet highly similar genomic regions (Srivastava *et al*., 2020). The options –keepDuplicates and –-gcBias was used transcript quantification to account for the high sequence similarity between some of the duplicated genes from the salmonid genome duplication (Lien *et al*., 2016) and correct for fragment-level GC biases of aligned reads, respectively. To calculate mRNA expression at the gene-level from output files from Salmon the sum of the raw read counts and normalized gene expression values (transcripts per million) were done with the R package tximport (Soneson *et al*., 2015).

### Differential gene expression analyses between samples with different photoperiod history

We carried out three different analyses of differential gene expression using gill RNAseq data to define a set of core genes whose expression during PST in the gill consistently depends upon photoperiodic history in the FW phase. Two of these analyses used previously published datasets (Iversen *et al*., 2020, 2021)(experiments 1 and 2) and the third analysis used data from this study (experiment 3) (Figure S1). In experiment 1 (Figure S1A) we contrasted 7 months old salmon (reared in fresh water) either raised on continuous light (LL group, n=6) or exposed to 60 days of short days (8L:16D) followed by 50 days in LL (SPLL group, n=6). In experiment 2 (Figure S1B) we contrasted 11 months old salmon (reared in fresh water) exposed to 2 weeks of short photoperiod (2w group, 8L:16D, n=6) with fish exposed to 8 weeks of short photoperiod (8w group, n=6). Both groups experienced an 8-week period on continuous light after the short photoperiod. In experiment 3 (Figure S1C) from this study we contrasted smolt raised on continuous light (LL, n=981) with smolt exposed to 6 weeks of short photoperiod treatments (8L:16D, n=927).

Differential expression in experiment 1 and 2 were analysed with EdgeR v3.42.4. Genes with low expression were filtered (min number of reads summed across all fish per group was 10) and normalized using trimmed means of M-values. A quasi-likelihood negative binomial generalized log-linear model with 0 as intercept was fitted to the data. Differentially expressed genes (DEGs) were identified in pairwise contrasts using empirical Bayes F-tests. P-values were adjusted for multiple testing using the BH method. Since large sample numbers have been shown to exaggerate false positive rates when applying standard differential gene expression methods (Li *et al*., 2022), DEGs from experiment 3 were identified using Wilcoxon rank-sum test. False discovery rate thresholds were defined as the square root of the minimum sample size per group, resulting in adjusted p-value cutoffs of <= 0.05 for experiments 1-2 and <=0.002 for experiment 3. Finally, upset plots of shared significant DEGs were made with the R package upsetR (Conway *et al*., 2017) to define a core set of photoperiodic history dependent smolt development genes identified as DEGs in all three analyses.

### Differential gene expression between fish with different SW performance

DEG analyses was also used to identify gene expression differences in gill samples from FW between smolts with different growth and mortality outcomes in SW. First, we contrasted gene expression in fish from the LL group which died within 40 days post seawater transfer versus fish that were still alive in November, approximately five months after sea transfer (i.e. excluding fish that died of disease or was otherwise lost). Second, we contrasted gene expression differences between loser fish with extremely low growth and ‘winners’ (i.e. fish with normal growth r). Classification of “losers” was based on the growth rate (i.e. gained weight divided by initial weight) where the threshold of 0.5 was decided based on the minimum between the two peaks in the growth rate distribution (see Figure 2A). Read counts were normalized across samples using EdgeR’s TMM method and then converted to Counts Per Million (CPM). These values were subsequently log-transformed to log2(CPM+1). Genes with zero reads in more than 90% of the samples were excluded, resulting in the removal of approximately 9,000 out of 47,000 genes. The final dataset of log-transformed expression values was used for Wilcoxon rank-sum tests. Fold change values were calculated as the difference of the mean log transformed expression values of the contrasted samples.

### Random forest prediction

To build prediction models for SW survival and growth using FW gill gene expression data from this study (experiment 3) we used the RandomForestClassifier from the sklearn python module (Pedregosa *et al*., 2011) with parameters n_estimators=1000 and class_weight=‘balanced’ (otherwise defaults). We included the log transformed gene expression as well as the commonly used smolt marker NKA ratio as features. NKA ratio was calculated by subtracting the log transformed gene expression of NKA α1a (ENSSSAG00000088099) from NKA α1b (ENSSSAG00000041746). This is equivalent to dividing α1b/α1a in the non-transformed expression values. Samples were randomly split into 80% training samples and 20% test samples. This was repeated 100 times, i.e. random split and model training, recording the importance scores of the genes and the prediction probabilities of each sample. The mean of the importance scores and the prediction probabilities of the test samples are reported in the analyses. We performed RF analyses for both ‘early death’ and ‘loser fish’ outcomes. Both early death and loser fish was defined as outlined for the DEG analyses.

### Identification of Gill Cell-Type Specific Genes Using Single Nuclei RNAseq

To identify genes specific to different cell types in gill tissues, we utilized the single nuclei RNA sequencing (snRNAseq) data provided by West et al. (2021). This dataset comprises snRNAseq analyses performed on gill tissues sampled at various stages of smoltification under different photoperiod regimes. Nuclei were assigned to 20 clusters based on their expression patterns using Seurat (Hao *et al*., 2024). Each cluster was subsequently annotated with a cell type using known marker genes and gene ontology (GO) analysis.

The expression levels of 2,000 variable genes were averaged across the nuclei in each cell-type cluster and each genes was assigned to the assigned cell type of the cluster with the highest average expression level. Since the original gene annotations from West et al. (2021) were based on NCBI identifiers, we converted these to Ensembl gene IDs using the g:Profiler’s g:Convert utility (Kolberg *et al*., 2023). This conversion resulted in 1,354 cell-type specific Ensembl genes mapped from 1,271 NCBI genes (i.e. not all genes could be mapped and some had multiple matches).

### Cell type enrichment analysis

Fisher’s exact test using the *fisher.test()* in R (R Core Team, 2021)was used to assess whether the differentially expressed genes (DEGs) in early death are associated with certain cell types. DEGs were identified based on their importance scores, with those exceeding 5e-5 (top ∼10%) being classified as DEGs. These DEGs were further divided into up-regulated or down-regulated based on the fold change. For each cell type, we evaluated the independence between a gene’s classification as a DEG and its association with that particular cell type. An odds ratio greater than one from this test suggests an enrichment of DEGs that are specifically expressed in that cell type. Specifically, for up-regulated DEGs, it suggests a higher abundance of the corresponding cell type in the gill samples from fish that died early, and conversely, for down-regulated DEGs, it indicates a greater abundance in the surviving fish. The p-values were adjusted for multiple comparisons using the *p.adjust()* function in R (R Core Team 2021) with the Benjamini & Hochberg.

## Results

### Defining a core set of photoperiodic history dependent smolt gill development genes

Our reanalysis of the datasets from Iversen et al. (2020, 2021) (experiments 1 and 2) identified 781 and 6,923 genes with significant different expression levels between gill samples from fish with different photoperiodic history during the PST (Figure S1D). DEG analyses between gills from fish with different photoperiodic histories in this study (i.e. LL *vs* 8L:16D and LL *vs* 12L:12D) identified a total of 14,209 genes which were DEGs in one or both contrasts (Figure S1D, Table S1). Finally, we intersected these three analyses to define a core set of 219 genes whose expression during PST gill consistently depends upon photoperiodic history in the FW phase (Table S2).

### Random forest prediction models for mortality and growth stagnation

The main aim of this study was to test if smolt gill gene expression profiles in fresh water could be used to predict seawater performance at the individual fish level, and secondarily to determine whether candidate markers defined by their sensitivity to photoperiodic history constituted useful SW performance predictors. We made RF prediction models for two seawater outcomes; early mortality (<40 days after seawater transfer) and stagnation of growth, referred to as loser fish.

Early mortality was substantial (>8%, Figure 1A) in the LL group, while extremely low in both groups of fish exposed to reduced photoperiods during the FW phase (12L:12D, n=8 fish = 0.8% mortality) and (8L:16D, n=6 fish = 0.6% mortality). Hence, we only used the LL group to develop the RF prediction model for early mortality.

**Figure 1:**
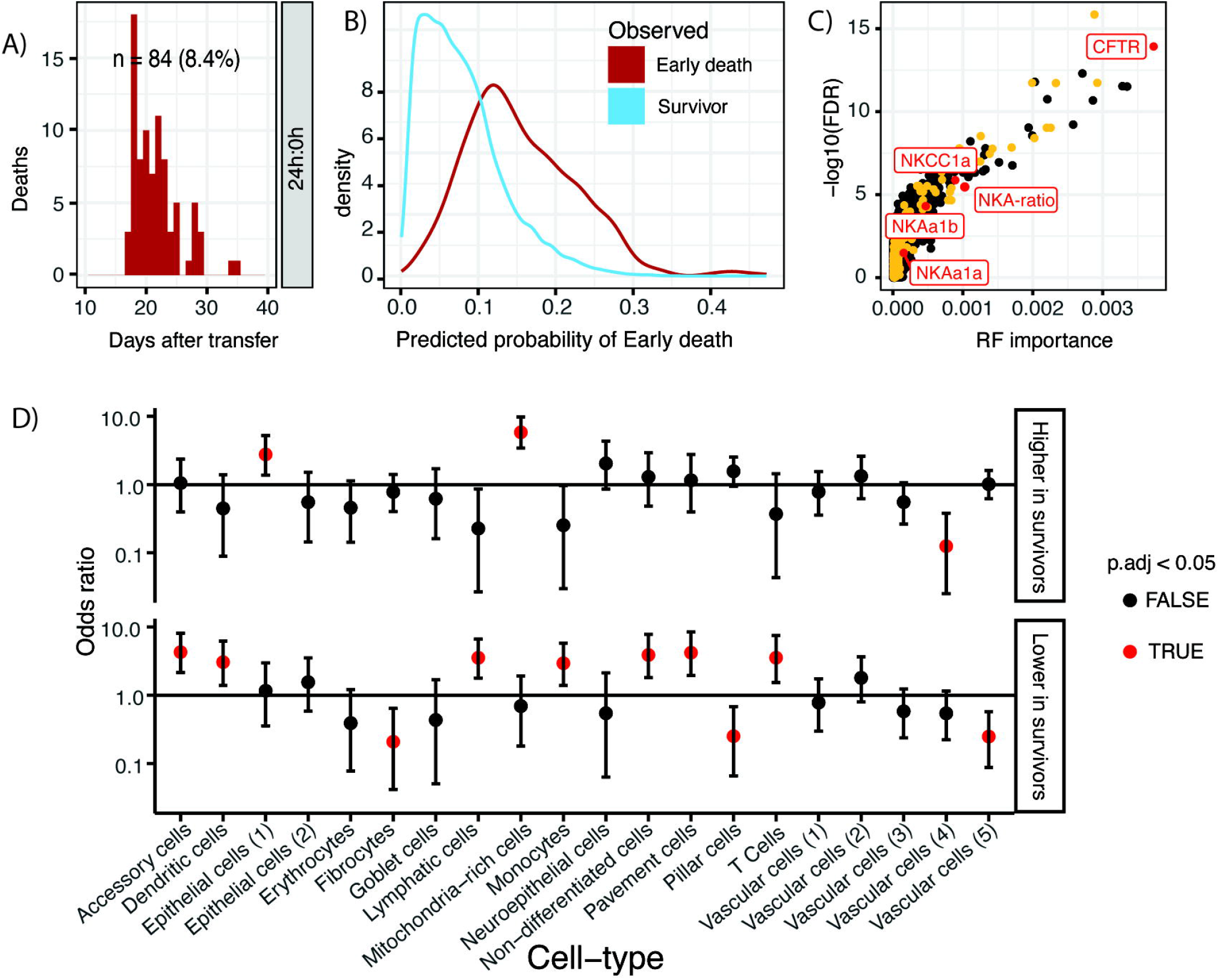
Random forest predictions of early mortality in seawater. A) Early mortality cases in LL group. B) Random forest model predictions for individual fish probability for mortality in an early phase (<40 days) following seawater transfer. Predictions are based on gill tissue gene expression profiles from smolts in fresh water prior to seawater transfer. C) Scatterplot of random forest model importance scores *vs* differential gene expression adjusted p-values from a contrast between fish dying early after seawater transfer and surviving fish. Yellow coloured dots represents genes with photoperiod history dependent regulation. Red dots represents classical smolt physiology states associated genes. D) Fisher test odds-ratios (i.e. enrichment) for cell type biassed/specific genes which are expressed higher and lower in surviving fish compared to fish dying early (<40 days) after seawater transfer.

**Figure 2.**
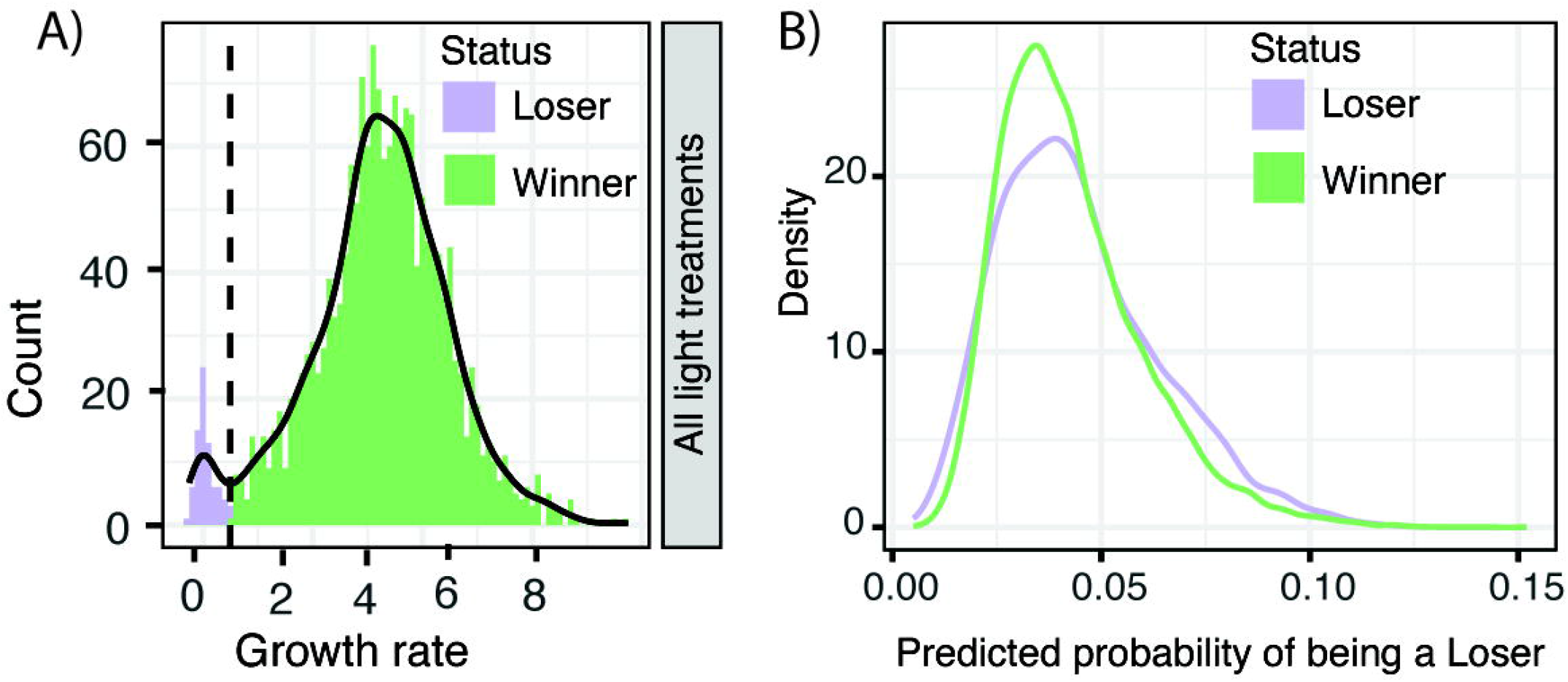
Random forest predictions of loser fish status in seawater. A) Classification of loser fish and winner fish based on growth rates in seawater. B) Random forest model predictions for individual fish classification of loser fish (low or no growth in seawater) and winner fish (normal growth rates) following seawater transfer. F) Scatterplot of random forest model importance scores *vs* differential gene expression adjusted p-values from a contrast between fish classified as loser fish (low or no growth in seawater) and winner fish (normal growth rates) following seawater transfer.

We tested the RF model on random subsets of LL fish not used in model training. This demonstrated good prediction capability (Figure 1B) with an average area under the receiver operating characteristic (auROC) of 0.82. In other words, in 82% of the predictions the model will correctly assign a higher absolute risk of mortality to a randomly selected fish that will die compared to a randomly selected fish which will not die.

Although an auROC > 0.80 is considered to be clinical useful in medicine (Çorbacıoğlu & Aksel, 2023), this prediction model will have relatively high false positive rate. Nevertheless, a ranked list of the importance scores of the RF model features (i.e. gene expression levels) can also be used to gain insights into biological processes which contribute to SW performance. We therefore performed an in-depth analysis of the gene importance scores. Interestingly, the most used molecular marker for smolt status, the two individual genes forming the NKA-subunits (NKAα1a and NKAα1b) had low importance scores in the prediction model (Figure 1C). We also included a proxy for the commercially available SmoltVision test (Pharmaq Analytic, Norway) which is the ratio of mRNA levels for the two NKA-subunits (NKAα1b:NKAα1a). Surprisingly, this NKA-ratio feature was only the 35^th^ most important feature in the RF model (Figure 1C, Table S3). However, another well-known smolt-gill gene, the cystic fibrosis transmembrane regulator (CFTR) (Hiroi & McCormick, 2012), had the highest importance in our RF model (Figure 1C). In addition, the top predictive genes (importance > 0.001, 37 genes, Table S3) also contained other genes linked to ion-transport such as the sodium/potassium-transporting ATPase subunit alpha-3 and the patj gene, which is involved in tight junction formation in mammals (Shin *et al*., 2005).

We then explored the predictive importance of genes with photoperiod-history dependent regulation (intersection between DEGs in experiment 1, 2, and this study, see Figure S1) for initial SW survival. Among the 219 photoperiodic history-dependent genes, 48% were in the top 10% most important genes in the RF model (yellow coloured points in Figure 1C). This was a significantly higher percentage than expected by chance (Fisher test p-value=9.53 × 10^−47^).

The development of the smolt gill involves a pronounced remodelling of its cellular composition, notably in terms of its complement of chloride cells and of immune cell types (McCormick, 2013; West *et al*., 2021). Leveraging recently published data on gill cell type-specific markers (West et al. 2021), we linked gene expression differences between surviving fish and those which died early after seawater transfer to the cell types these differentially expressed genes were associated with. Genes associated with mitochondria-rich cells and one subtype of epithelial cells (1) were significantly more likely (FDR<0.05) to be expressed more strongly in survivors compared to fish dying within 40 days of SW transfer (Figure 1D). Genes associated with seven cell types, including three immune-system related cells, were significantly more likely (FDR<0.05) to be expressed at lower levels in survivors compared to fish that died within 40 days of SW transfer (Figure 1D).

Finally, we used a similar RF approach to predict loser fish status, which is a common challenge in aquaculture. Short photoperiod exposure protocols were not associated with loser fish status; hence we combined all loser fish in one group (Figure 2A). Our results clearly demonstrate that it is not possible to use gill gene expression in the FW phase to train an RNF model to predict loser fish status in SW (Figure 2B).

### Including external smolt characteristics in the RF model

External characteristics such as size, condition factor, and skin colouration are being used to evaluate smolt status and development in salmon aquaculture (Ytrestøyl *et al*., 2023). Although our study was focussing on transcriptome-based markers, we therefore also build an RF model including size and a smolt index score in June prior to sea water transfer.

Our results showed that body weight had the highest model importance score for early seawater mortality and that skin colouration had higher model importance than the commercially used NKA-ratio (Figure 3A). Condition factor ranked below the NKA-rato in model importance (Figure 3A). It is interesting to note that even though the LL group had ten times higher mortality in early sea water phase, the proportion of silvery coloured fish among fish that died were similar across all smolt production groups (Figure 3B).

**Figure 3.**
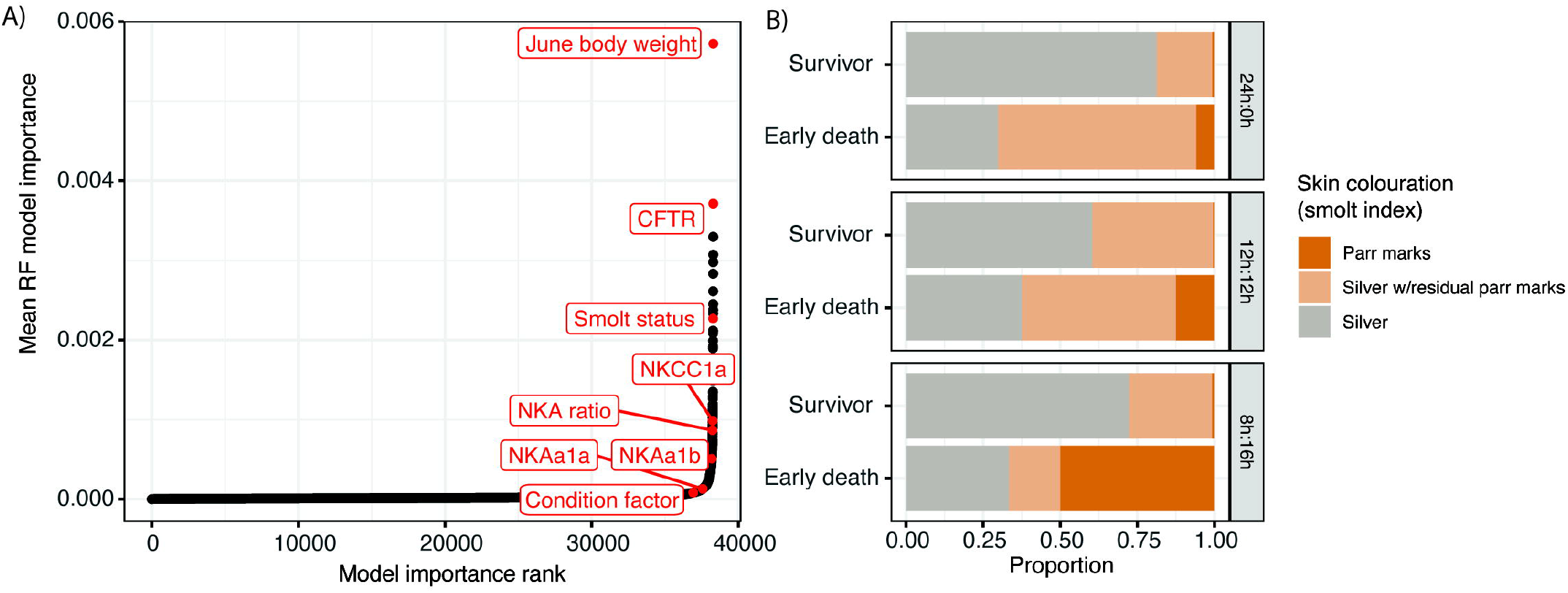
Random forest predictions including external smolt characteristics. A) Feature importance plotted for random forest model for early mortality in sea water. This model included three external smolt characteristics; body size, condition factor and skin colouration (i.e. smolt index). B) Proportions of smolt index scores in the three smolt groups reared under different photoperiod histories.

## Discussion

In the present study we aimed at identifying and validating gene expression markers for smolt status by using gill gene expression at the end the smolt rearing in FW to predict survival and performance in the SW phase. We could not predict loser fish outcomes based on FW gill transcriptomes, indicating that this phenomenon is not mechanistically linked to smolt gill development and ion regulation ability. Conversely, the prediction model for SW survival had reasonable discriminatory ability (auROC=0.84), and results from the RF model (Figure 2C) provide novel biological insights into the smolt gill and development of new biomarkers for smolt status.

The most common molecular biomarker for smolt status is the NKA-subunit gene expression ratio (NKAα1b: NKAα1a) (Reviewed in Takvam *et al*., 2024). Surprisingly, in this study (Figure 1C) the NKA-ratio was only ranked as the 35^th^ most important feature in the prediction model for early mortality in SW. This result is in line with a recent experiment by Kahw et al. (2021), which found large variation in NKA-ratio related to fish genetics and smolt production protocols (0+ vs 1+) and concluded that NKA-ratio was not a robust biomarker for smolt status. Hence, Kahw et al. (2021) and our findings in this study questions the reliability of the NKA-ratio as a good predictor for seawater survival and performance in aquaculture smolt production.

Our large-scale transcriptomic approach using gene expression profiles from individual fish therefore represents an important step towards identifying new and improved biomarkers for smolt status. One interesting aspect of our list of putative smolt biomarkers is the highly significant overrepresentation of core photoperiod-history sensitive genes (Fisher test, p-value = 9.529355e-47, Figure 1C). The most predictive gene CFTR (Figure 1C) was not in the core photoperiod-history sensitive gene set, however CFTR was deemed photoperiodic-history sensitive in two out of three experiments (Table S1). Among the top 36 features with highest prediction model importance (> 0.001), 12 were in the core photoperiod sensitive gene list, including calpain 2, egln2, and TEFa. Interestingly, calpain2 was ranked as the gene with the strongest photoperiod-history sensitive regulation during PST in Iversen et al. (2020). Egln2 is known to interact with hypoxia inducible factors (HIFs) in human studies according to the Molecular Interaction Database (https://www.ebi.ac.uk/intact/) and in zebrafish larvae there is a bidirectional crosstalk between HIFs and glucocorticoid signalling (Marchi *et al*., 2020). Furthermore, the D-element transcription factor TEFa, which is implicated in light - circadian interactions in zebrafish (Vatine *et al*., 2009), is a core part of photoperiod transduction in mammals (Dardente *et al*., 2010) and is directly responsive to cortisol in salmon (West *et al*., 2020). The presence of egln2 and TEFa among the most predictive genes thus suggests a glucocorticoid-signalling link to PST gill development and initial SW survival. In conclusion, our results points to a mechanistic link between poor smolt development in the LL group and dysregulation of molecular developmental processes that are naturally regulated by photoperiod signals. Focusing on these “winter signal” affected genes is therefore an exciting avenue in future research and validation of novel biomarkers for smolt physiology.

In the models including external features smolt size were ranked higher in model feature importance compared to any of the gene expression phenotypes (Figure 3A). This is not surprising as the ability to tolerate and survive osmotic stress increases with fish size, irrespective of smolt development (Bjerknes *et al*., 1992; Parry, 1960). Skin colouration and condition factor did however rank lower than many gene expression phenotypes in their model importance score, and the increased mortality in the LL group were not associated with lower proportion of fish with silvery skin colour (Figure 3B). This aligns with the notion that development of smolt physiology happens through a suite of parallel and independent processes, and that not all are causally linked to the molecular basis for sea water tolerance.

One caveat related to our prediction model results is that the data we used to build the model was limited to data from fish raised on LL. This was because initial survival was >99% in smolts raised using exposure to short photoperiods. We did however try to apply our model to the limited number of fish in 8:16 and 12:12 that suffered early sea water mortality, which showed that the fish with highest probability of early sea water mortality did in fact die (Supplementary Figure 2). However there was not a significant difference in the distribution of probability of mortality for the few fish that did die compared to the vast majority that survived in these groups. Even though an LL protocol can compromise the development of hypo-osmoregulatory ability as well as other smolt development characteristics, our results are still of high relevance to the industry due to extensive use of LL in smolt production.

## Author contributions

Mattis J. van Dalum: investigation, formal analysis, visualization, writing

Bjarne Gjerde: conceptualization, experimental design, manuscript editing

Solomon Boison: conceptualization, experimental design, manuscript editing

Lars Grønvold: investigation, formal analysis, writing, manuscript editing

David G. Hazlerigg: conceptualization, funding acquisition, writing and editing, supervision

Even Jørgensen: fish rearing and sampling, manuscript editing

Turid Mørkøre: fish phenotype recording, manuscript editing

Simen R. Sandve: conceptualization, funding acquisition, writing and editing, supervision

Anja Striberny: fish rearing and sampling, writing

Domniki Manousi: bioinformatic data analyses, manuscript editing

Trine Ytrestøyl: fish phenotype recording, manuscript editing

## Acknowledgements

Thanks to MOWI for providing fish for this experiment and Sofie Robertson for exploring the use of RF prediction models using gene expression data.

## Funding acknowledgements

This study was funded by the Norwegian Seafood Research Fund (project no. 901589).

## Data availability

All the reads reads from transcriptome data from the new experiments presented in this paper is deposited at ENA under the project number ERP131717. All scripts and data to reproduce figures and statistical tests can is deposited at zenodo (https://doi.org/10.5281/zenodo.14177709)

## Figure and table legends

**Figure S1.Defining a core set of photoperiod history-sensitive genes**

A) Experiment 1: a ‘short day’ experiment comparing the effect of FW rearing in continuous light (LL) throughout with exposure to 8-h light / 24-h for a 6 week period. B) Experiment 2: a ‘winter length‘ experiment in which the duration of exposure to 8-h light / 24-h was either 2-or 8 weeks. C) Experiment 3: data from this study in which the effect of rearing on LL throughout were compared with exposure to either 12-h or 8-h light / 24-h for 6 weeks. D) Upset plot showing differentially expressed genes (DEGs) in each of the experiments. DEGs were filtered down to a common set of genes that had photoperiod sensitive regulation in all three experiments (FDR adjusted p-value <= 0.002 for experiment 3 and <=0.05 for experiments 1, and 2).

**Figure S2. Random forest predictions of early mortality applied on 12:12 and 8:16 fish.**

Predicted probabilities for early mortality in sea water for fish reared under 12:12 and 8:16 regimes. Note that the random forest model is trained on the fish from the LL group.

**Table S1. Results from all DEG analyses and RF model**

**Table S2. Core photoperiod-history sensitive genes**

**Table S3. Top predictive genes for mortality in the initial SW phase**

