## Supplementary figures and images for "Transcriptomic profiling of gill biopsies to define predictive markers for seawater survival in farmed Atlantic salmon"

### Figure1S.tif

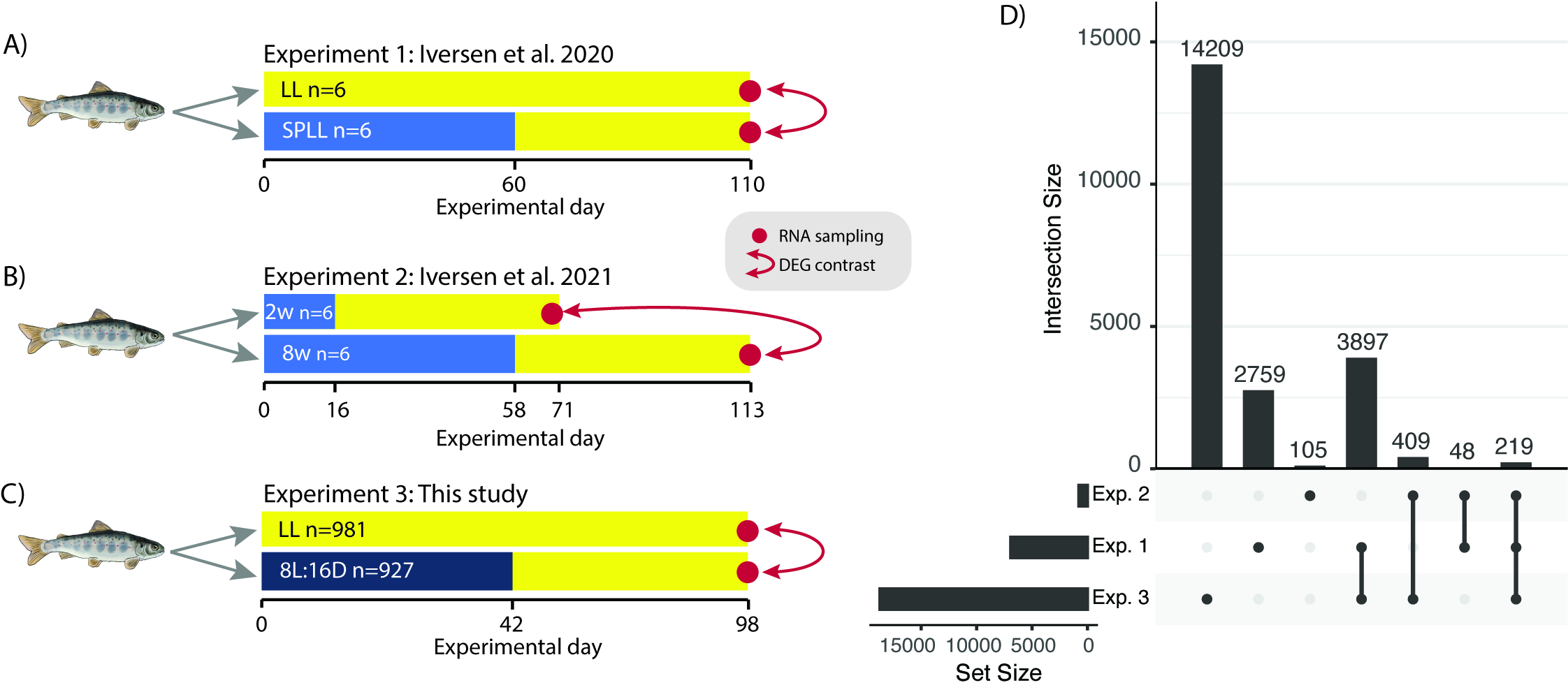

### Figure2S.tif

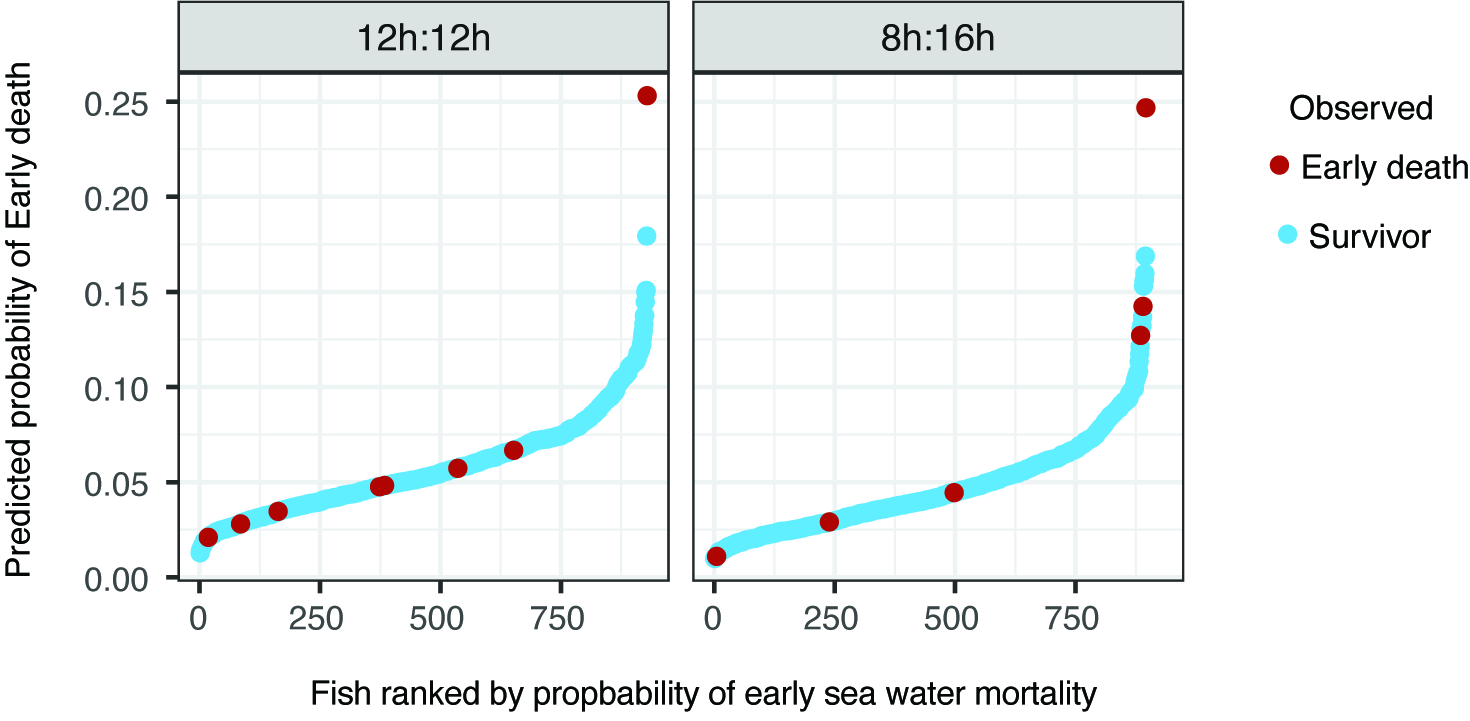
